# Molecular basis of functional compatibility between ezrin and other actin-membrane associated proteins during cytokinesis

**DOI:** 10.1101/2020.11.30.403253

**Authors:** Guang Yang, Shota Hiruma, Akira Kitamura, Masataka Kinjo, Mithilesh Mishra, Ryota Uehara

**Author notes:** Corresponding author: Ryota Uehara, Faculty of Advanced Life Science, Hokkaido University, Kita 21, Nishi 11, Kita-ku, Sapporo 001-0021, Japan.

## Abstract

The mechanism that mediates the interaction between the contractile ring and the plasma membrane during cytokinesis remains elusive. We previously found that ERM (Ezrin/Radixin/Moesin) proteins, which usually mediate cellular pole contraction, become over-accumulated at the cell equator and support furrow ingression upon the loss of other actin-membrane associated proteins, anillin and supervillin. In this study, we addressed the molecular basis of the semi-compatibility between ezrin and other actin-membrane associated proteins in mediating cortical contraction during cytokinesis. We found that depletion of supervillin and anillin caused over-accumulation of the membrane-associated FERM domain and actin-binding C-terminal domain (C-term) of ezrin at the cleavage furrow, respectively. This finding suggests that ezrin differentially shares its binding sites with these proteins on the actin cytoskeleton or inner membrane surface. Using chimeric mutants, we found that ezrin C-term, but not the FERM domain, can substitute for the corresponding anillin domains in cytokinesis and cell proliferation. On the other hand, either the membrane-associated or the actin/myosin-binding domains of anillin could not substitute for the corresponding ezrin domains in controlling cortical blebbing at the cell poles. Our results highlight specific designs of actin- or membrane-associated moieties of different actin-membrane associated proteins with limited compatibility, which enables them to support diverse cortical activities on the shared actin-membrane interface during cytokinesis.

## Introduction

Contractile force generated by the actin cytoskeleton-based contractile ring drives cell membrane deformation in cytokinesis. In this process, different types of proteins containing both actin- and membrane-associated domains concentrate at the cleavage furrow that forms at the cell equator. Anillin consists of the N-terminal actin/myosin-binding domains and the C-terminal phosphatidylinositol 4,5-bisphosphate (PIP2)-binding domain, through which it anchors the contractile ring to the cell equator for membrane deformation in different animal cells [1–3]. Supervillin also possesses actin- and myosin-binding domains, associates with the plasma membrane, and mediates cleavage furrow ingression in mammalian cells [4–6]. ERM proteins associate with the membrane through their conserved N-terminal FERM domain, bind to F-actin through the C-terminal domain, and accumulate prominently at the cleavage furrow [7–10]. Depletion of anillin and supervillin causes severe and modest furrowing defects, respectively, during cytokinesis [6, 11–13]. On the other hand, depletion of ERM proteins does not affect cytokinesis progression either in human or *Drosophila* cells [10, 14, 15]. ERM depletion instead affects the cell polar cortex’s reorganization during cytokinesis, particularly perturbing membrane retraction during cortical blebbing [14–16]. Moreover, deregulation of moesin’s phosphorylation cycles impairs cortical dynamics at the cell poles, blocks normal cell elongation during anaphase, and causes cytokinesis defects [17, 18]. However, it remains mostly elusive why these actin-membrane associated proteins are different in their functions and relative contributions to cytokinesis control, despite the apparent similarity in their molecular design.

A possible reason for the difference in these actin-membrane associated proteins’ functionality may be the distinctive characteristics of their molecular domains. For example, a myosin-binding domain is closely located beside the actin-binding domain in anillin and supervillin but is absent in ERM proteins, which may limit the interaction of ERM proteins with the contractile ring [2, 5]. Moreover, anillin’s actin-binding domain contains characteristic multiple actin-binding sites, enabling F-actin bundling with the distinctive polymorphic modes [19, 20]. Such distinctive properties of molecular domains may promote specialization of their roles in reorganizing the actin-membrane interface at the cleavage furrow. Moreover, differences in their interactomes would also enable the diverse functions of these proteins. For example, anillin interacts with an essential cytokinesis regulator RhoA through its C-terminal Rho-binding domain, facilitating the importin-mediated recruitment of anillin and retention of RhoA at the cortex to ensure the transduction of RhoA signaling [13, 21, 22]. Supervillin interacts with a subset of cytokinesis-relevant proteins, including myosin light chain kinase, PRC1, and EPLIN, potentially promoting myosin activation at the furrow [6, 12, 23, 24]. Recent studies revealed that ezrin interacts with novel cytokinesis-relevant proteins CLIC1/4, and that ezrin and CLIC1/4 mutually support their localization to the cleavage furrow [25, 26]. In *Drosophila* cells, a subset of cortically-localized moesin directly interacts with microtubules for regulating cortical rigidity during pre-anaphase and cytokinesis [27]. Besides the interaction with cytokinesis-relevant proteins, ezrin also interacts with a Rho-activating factor MYOGEF through the FERM domain and mediates its recruitment to the membrane blebs, which is required for RhoA activation at the blebs [28]. Such distinctive protein interactions would give unique roles to these actin-membrane associated proteins, potentially making them less compatible with each other in supporting cortical dynamics during cytokinesis.

Interestingly, however, we previously found that, when we depleted anillin and supervillin, ezrin was over-accumulated at the cleavage furrow in HeLa cells. In this condition, ERM proteins became engaged in furrow ingression, whereas they were dispensable for furrow ingression in the presence of anillin and supervillin [15]. This compensatory over-accumulation of ezrin indicates semi-compatibility between ezrin and the other actin-membrane associated proteins. However, it remains unknown which molecular domains of these actin-membrane associated proteins support or limit their compatibility in cytokinesis control. In this study, by analyzing the localization, dynamics, and function of ezrin molecular domains, we investigated the mechanism that determines both the unique and shared roles of ERM proteins in controlling cortical contractile activity during cytokinesis in human cells.

## Materials and methods

### Cell culture

HeLa-Kyoto cell lines were cultured in Dulbecco’s modified Eagle’s medium (DMEM, Wako, Japan) supplemented with 10% fetal bovine serum (FBS) and 1× antibiotic-antimycotic (Sigma-Aldrich, St. Louis, MO). For live imaging, cells were cultured in phenol red-free DMEM (Wako) supplemented with 10% FBS and 1× antibiotic-antimycotic on cover glass-bottom culture dishes (P35G-1.5-14-C, Mattek, Ashland, MA).

### siRNA, plasmid, and nucleotide transfection

The siRNAs used in this study are 5’-CGAUGCCUCUUUGAAUAAAtt-3’ (anillin#1) [13], 5’-AGCTTACAGACTTAGCATAtt-3’ (anillin#2; targeting 3’-UTR sequence), 5’-GAGAACAAGGGAAUGUUGAGAGAat-3’ (supervillin) [15], 5’-CGUGGGAUGCUCAAAGAUAtt-3’ (ezrin) [15], 5’-GGCTGAAACTCAATAAGAAtt-3’ (moesin) [15], 5’-GGAAGAACGTGTAACCGAAtt-3’ (radixin) [15], and 5’-CGUACGCGGAAUACUUCGAtt-3’ (luciferase; DNA is shown in lowercase) [15]. siRNA transfection was performed using Lipofectamine RNAiMAX (Thermo Fisher Scientific, Waltham, MA). The plasmid vectors constructed in this study are listed in Table S1. DNA transfection was performed using JetPEI (Polyplus-transfection, Illkirch, France). HeLa cell lines stably expressing the GFP-tagged ezrin or anillin mutant genes were obtained by selecting GFP-positive cells in the presence of 500 μg/mL G418. Anillin-GFP or RFP-α-tubulin stable line has been previously described [29].

### Cell fixation

For the cell fixation, cells were fixed with 3.2% paraformaldehyde in phosphate-buffered saline [PBS] for 10 min and permeabilized with 0.5% Triton-X100 in PBS supplemented with 0.1 M glycine [GPBS] for 10 min at 25°C.

### Immunoblotting

For immunoblotting, proteins separated by SDS-PAGE were transferred on to Immun-Blot PVDF membrane (Bio-Rad, Hercules, CA). The blotted membranes were blocked with 0.3% skim milk in TTBS (50 mM Tris, 138 mM NaCl, 2.7 mM KCl, and 0.1% Tween 20), incubated with the primary antibodies overnight at 4°C or for 1h at 37°C, and incubated with the secondary antibodies for 30 min at 37°C. Each step was followed by 3 washes with TTBS. For signal detection, the ezWestLumi plus ECL Substrate (ATTO, Tokyo, Japan) and a LuminoGraph II chemiluminescent imaging system (ATTO) were used.

### Antibodies

Mouse anti-GAPDH (sc-32233, Santa Cruz Biotechnology, Dallas, TX; 1:100), mouse anti-GFP (mFX75, Wako; 1:500), mouse anti-β-tubulin (10G10, Wako; 1:1000), rabbit anti-supervillin (NBP1-90363, Novus Biotechnologicals, Littleton, CO; 1:500), mouse anti-anillin #1 (sc-271814, Santa Cruz Biotechnology; 1:100), goat anti-anillin #2 (sc-54859, Santa Cruz Biotechnology, Dallas, TX; 1:100), mouse anti-ezrin (sc-58758, Santa Cruz Biotechnology; 1:100), rabbit anti-radixin (EP1862Y, GeneTex, Irvine, CA; 1:10000), rabbit anti-moesin (#3150, Cell Signaling Technology, Danvers, MA; 1:1000), and horseradish peroxidase-conjugated secondary antibodies (Jackson ImmunoResearch Laboratories, West Grove, PA; 1:1000) were purchased from the suppliers and used at the dilutions as indicated.

### Cell imaging

For fixed cell imaging, cells were observed under a TE2000 microscope (Nikon, Japan) equipped with a ×60 1.4 NA Plan-Apochromatic, a CSU-X1 confocal unit (Yokogawa, Tokyo, Japan), and an iXon3 electron multiplier-charge coupled device (EMCCD) camera (Andor, Belfast, United Kingdom) or ORCA-ER CCD camera (Hamamatsu Photonics, Hamamatsu, Japan), or a Ti-2 microscope (Nikon) equipped with ×60 1.4 NA Apochromatic, and Zyla4.2 sCMOS camera (Andor). Image acquisition was controlled by μManager (Open Imaging). For quantification of fluorescence intensity of EGFP-tagged ezrin mutants at the cleavage furrow and the polar cortex, dividing cells with their furrow width ranging from 2.5 to 10 μm (Fig. 2E) or 4 to 13 μm (Fig. 3C) were analyzed. Line profiles were obtained using 10 or 20-pixel wide lines across and along the cell division axis (for fluorescence measurement at the furrow and the poles, respectively) in Image J software. Then, fluorescence intensity values at the points corresponding to the cleavage furrow or the polar cortex in the line profiles were subtracted by background fluorescence intensity outside the cells and subjected to furrow/pole ratio calculation. The frequency of abnormally large bleb formation in Fig. 6 was quantified by counting the number of the events taking place during cytokinesis (defined as the duration from anaphase onset to the time point when furrow width reached less than 4 μm) and dividing the event number by the duration of cytokinesis. We defined the blebs whose maximum area size exceeded 40 μm^2^ as abnormally large blebs. The frequency of multinucleated cells was counted using the cell counter plugin in Image J.

### Fluorescence recovery after photobleaching (FRAP)

We performed FRAP experiments for ezrin-EGFP using an LSM510 microscope (Carl Zeiss, Jena, Germany) equipped with a C-Apochromat ×40 1.2 NA W Corr. UV-VIS-IR water immersion objective (Carl Zeiss). Images were acquired every 1 s. Photobleaching was conducted at 3 μm-diameter circle region at the cleavage furrow using 488 nm laser (88 μW for 2.08 s) after 2 frames of pre-bleaching imaging. Dividing cells with their furrow width ranging from 7 to 21 μm (at the first time frame of pre-bleaching imaging) were analyzed. Normalized fluorescence intensity at the bleached furrow region was obtained by dividing fluorescence intensity at each time point after photobleaching by pre-bleaching fluorescence intensity. We measured fluorescence intensity at the furrow region using round-shaped regions of interest (ROIs) with diameters of 2.1 μm in ImageJ software. FRAP curves were fitted using a single exponential equation;

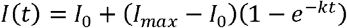

where *I*_*0*_ or *I*_*max*_ is fluorescence intensity immediately after photobleaching or plateau fluorescence intensity after recovery, respectively, and *k* is fluorescence recovery rate constant. Non-linear curve fitting was conducted using the solver add-in of Excel software (Microsoft). Half recovery time τ_1/2_ is then obtained using equation;

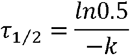

### Statistical analysis

Analyses for significant differences among different samples were conducted using the two-tailed Student’s *t*-test. Statistical significance was set at P < 0.05.

### Colorimetric cell proliferation assay

For cell viability assay, 4,000 HeLa WT or ezrin-C-term-EGFP-anillin-C-term cells were seeded on each well of 24-well plates. Immediately after the cell seeding, cells were transfected with siRNA targeting luciferase (for mock depletion) or anillin 3’-UTR. Then, RNAi treatment was repeated every 24 h. Ninty-six h after the cell seeding, 5% Cell Counting Kit-8 (Dojindo) was added to the culture, incubated for 4 h, and absorbance at 450 nm was measured using the Sunrise plate reader (Tecan). The absorbances of anillin-depleted samples were normalized to those of the corresponding mock-depleted controls.

## Results

### Compensatory accumulation of ezrin takes place without changing its molecular turnover dynamics

To gain insight into how ezrin over-accumulates at the cleavage furrow upon depleting other actin-membrane associated proteins [15], we compared molecular turnover dynamics of ezrin-EGFP at the cleavage furrow in control, supervillin-, or anillin-depleted HeLa cells using FRAP (Fig. 1A-D). The accumulation of ezrin-EGFP at the furrow increased upon depletion of supervillin or anillin, which was indicated by the significant increase in the equatorial to polar cortical fluorescence signal ratio (Fig. 2A, B, E, and 3A-C). We reasoned that if supervillin or anillin influences ezrin localization through direct modulation of its association with the cleavage furrow, the turnover of ezrin-EGFP at the furrow would change upon depletion of these proteins. In control cells, FRAP of ezrin-EGFP at the cleavage furrow took place with an estimated half recovery time of 29 ± 1.8 s (mean ± s.e., n=68 from four independent experiments; Fig. 1B-D). The FRAP profile of ezrin-EGFP at the cleavage furrow was similar to those previously reported at the cell cortex in interphase cells [30]. Depletion of supervillin or anillin did not significantly change the turnover of ezrin-EGFP at the cleavage furrow (an estimated half time recovery of 32 ± 2.5 s or 30 ± 2.5 s, respectively, n=51 or 75 from four independent experiments, respectively; Fig. 1B-D). These results suggest that supervillin and anillin are not involved in modulating ezrin dynamics at the furrow.

**Fig. 1:**
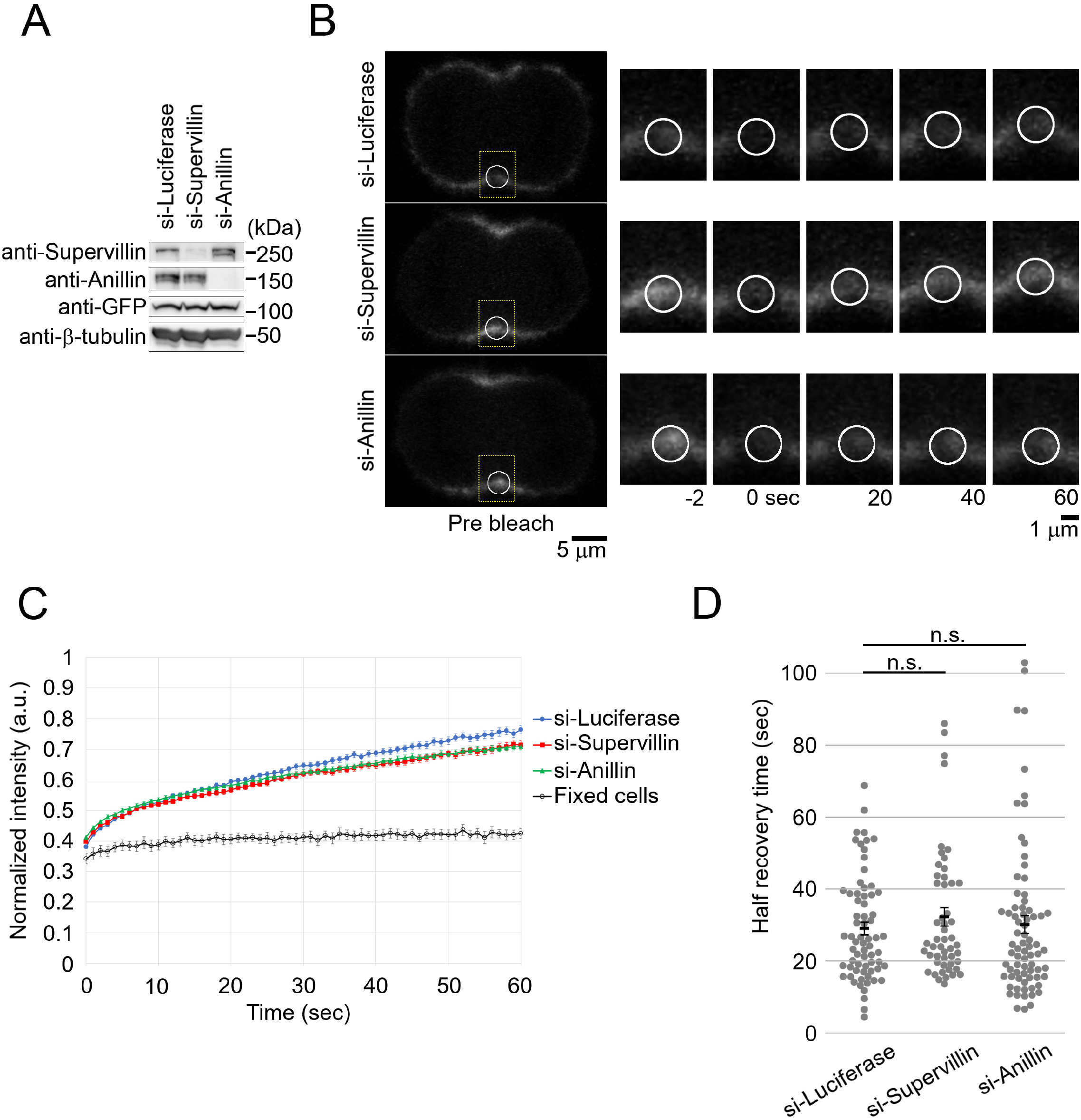
Depletion of supervillin does not change the molecular turnover dynamics of ezrin-EGFP at the cleavage furrow in HeLa cells. (**A**) Immunoblotting of GFP, supervillin, and anillin in RNAi-treated HeLa cells expressing ezrin-EGFP. β-tubulin was detected as a loading control. (**B**) Live images of RNAi-treated cells expressing ezrin-EGFP before and after photobleaching. Photobleached regions at the cleavage furrow are indicated by open circles in the left panels. Boxed regions at the cleavage furrow are enlarged in the right panels. Regions of interest (ROIs) used for intensity analysis are indicated by open circles in the right panels. Photobleaching was conducted at 0 s. (**C**, **D**) Quantification of fluorescence recovery after photobleaching in B (C), and estimated half fluorescence recovery time (D). Means ± standard errors (SE) of normalized fluorescence intensity at the cleavage furrow taken from at least 51 cells from four independent experiments (live cells) or 10 cells from three experiments (fixed cells). In live cell analysis, there was no statistically significant difference between control and the other two samples (*p* = 0.27 or 0.64 for supervillin- or anillin-depleted cells, respectively, two-tailed *t*-test).

**Fig. 2:**
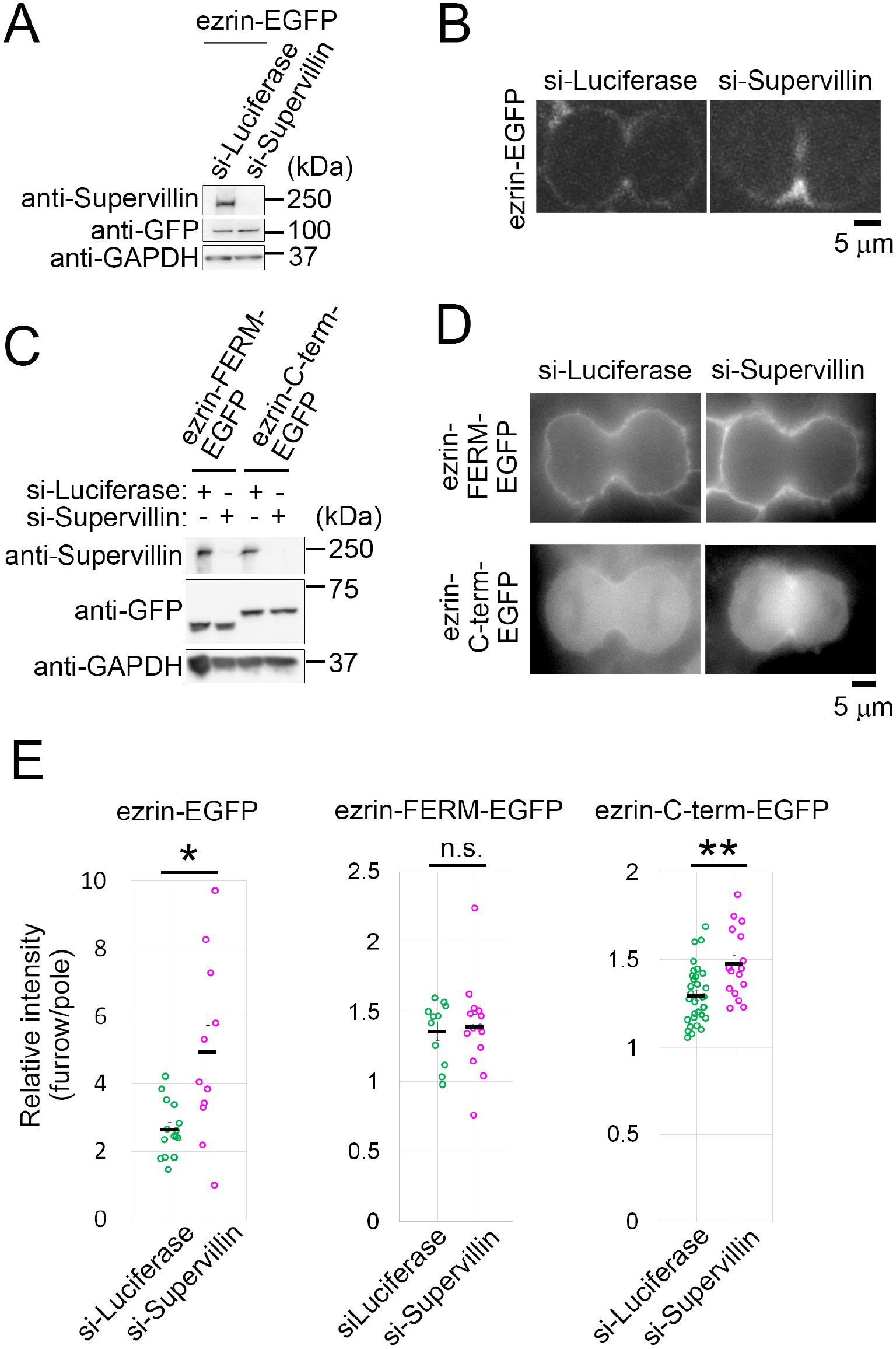
Localization of ezrin-FERM-EGFP and ezrin-C-term-EGFP at the cleavage furrow in control and supervillin-depleted HeLa cells. (**A**, **C**) Immunoblotting of GFP and supervillin in mock- or supervillin-depleted HeLa cells expressing EGFP-tagged full-length ezrin (A) or ezrin truncates (C). GAPDH was detected as a loading control. (**B**, **D**) Fluorescent microscopy of EGFP-tagged full-length ezrin (B) or ezrin truncates (D) in mock- or supervillin-depleted cells. (**E**) Quantification of cleavage furrow/polar cortex ratio of fluorescence signals in B and D. Means ± SE of at least 11 cells from two independent experiments for each condition. Asterisk indicates significant difference from control (* *p* < 0.05, ** p < 0.01, two-tailed *t*-test).

**Fig. 3:**
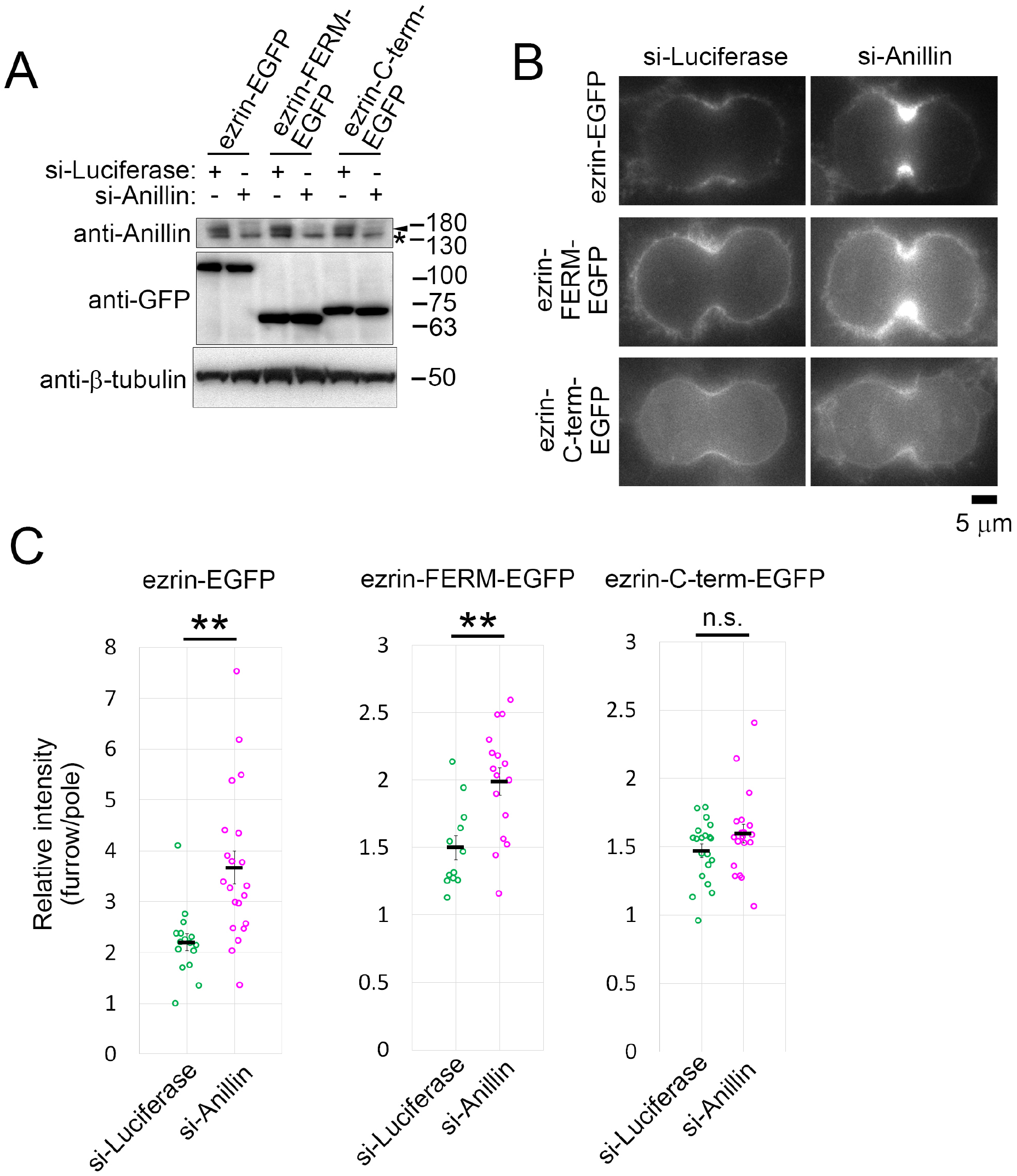
Localization of ezrin-FERM-EGFP and ezrin-C-term-EGFP at the cleavage furrow in control and anillin-depleted HeLa cells. (**A**) Immunoblotting of GFP and anillin in mock- or anillin-depleted HeLa cells expressing EGFP-tagged ezrin full length, FERM, or C-terminal domain. β-tubulin was detected as a loading control. The arrowheads indicate endogenous anillin and the asterisk indicates nonspecific bands. (**B**) Fluorescent microscopy of EGFP-tagged ezrin mutants in mock-, or anillin-depleted cells. (**C**) Quantification of cleavage furrow/polar cortex ratio of fluorescence signals in B. Means ± SE of at least 12 cells from two independent experiments for each condition. Asterisk indicates significant difference from control (** p < 0.01, two-tailed *t*-test).

### Anillin or supervillin depletion causes over-accumulation of different domains of ezrin at the cleavage furrow

All ERM proteins, anillin, and supervillin can associate with the actin cytoskeleton and membrane at the cleavage furrow. Therefore, anillin and supervillin may usually suppress ERM proteins’ accumulation through competition for shared binding sites either on the actin cytoskeleton or inner membrane surface. To test this idea, we investigated the effects of supervillin depletion on the accumulation of EGFP-tagged membrane-associated FERM domain of ezrin (ezrin-FERM-EGFP) or actin-binding C-terminal half of ezrin (ezrin-C-term-EGFP) at the furrow (Fig. 2A-E). The depletion of supervillin resulted in increased accumulation of ezrin-C-term-EGFP at the furrow but did not change that of ezrin-FERM-EGFP (Fig. 2E). We also tested the effect of anillin depletion on the accumulation of the ezrin truncates (Fig. 3A-C). In contrast to supervillin depletion, anillin depletion caused an increase in the accumulation of ezrin-FERM-EGFP at the furrow but not ezrin-C-term-EGFP (Fig. 3C). These results suggest that ezrin differentially shares its binding sites on membrane or actin cytoskeleton with anillin or supervillin, respectively, at the cleavage furrow.

### Compatibility of actin-binding domains between ezrin and anillin for the induction of furrow contraction

The above results indicate that ezrin potentially associates with the membrane-actin cytoskeleton interface relevant to furrow ingression activity. This prompted us to test whether the FERM domain and/or C-terminal actin-binding domain of ezrin can support furrowing activity when swapped with the corresponding original domains of anillin. For this, we constructed chimeric mutant genes containing the FERM domain with anillin N-terminal actin-/myosin-binding domains (ezrin-FERM-EGFP-anillin-MABD) or ezrin C-terminal actin-binding domain with anillin C-terminal membrane-binding domain (ezrin-C-term-EGFP-anillin-C-term; Fig. 4A and S1). The functionality of these chimeric proteins in furrow ingression was investigated by live imaging in HeLa cells in which endogenous anillin was depleted using 3’-UTR targeting siRNA (Fig. 4B; immunoblotting shown in Fig. S1). When transiently expressed in HeLa cells, ezrin-FERM-EGFP-anillin-MABD broadly distributed at the cellular cortex with weak concentration at the equator during cytokinesis (Fig. 4B). Ezrin-C-term-EGFP-anillin-C-term strongly accumulated at the cleavage furrow (Fig. 4B). Anillin depletion severely suppressed furrow ingression with frequent furrow regression, but an exogenous expression of EGFP-anillin substantially restored normal dynamics of furrow ingression in anillin-depleted cells (Fig. 4C and D). In contrast, transient expression of EGFP-anillin-MABD, EGFP-anillin-C-term, ezrin-EGFP, ezrin-FERM-EGFP, ezrin-C-term-EGFP, or ezrin-FERM-EGFP-anillin-MABD did not restore furrow ingression in anillin-depleted cells. Interestingly, however, the expression of ezrin-C-term-EGFP-anillin-C-term substantially restored furrowing activity in anillin-depleted cells (Fig. 4B-D). However, the equatorial cortex was frequently deformed with irregular waviness during furrow ingression in the anillin-depleted cells expressing ezrin-C-term-EGFP-anillin-C-term (the arrow in Fig. 4B and E), which was much less frequent in EGFP-anillin-expressing cells. These data demonstrate that the actin-binding domain, but not the membrane-associated domain of ezrin can substitute for anillin’s corresponding domain for furrowing activity.

**Fig. 4:**
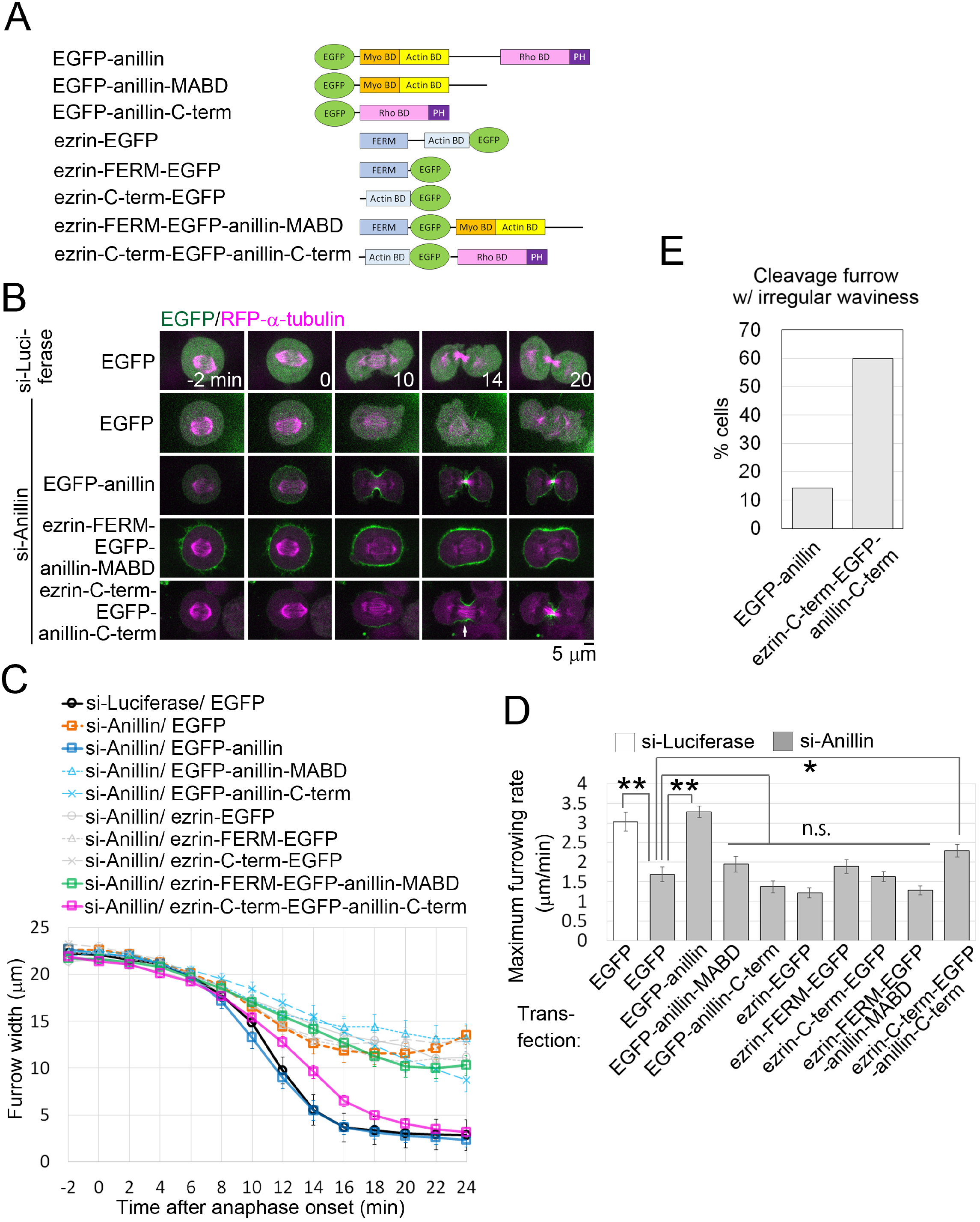
Ezrin C-terminal domain can substitute for anillin actin- and myosin-binding domains for supporting furrow ingression. (**A**) Schematic structure of truncated or chimera genes used in the experiments. (**B**) Live imaging of HeLa cells expressing EGFP-tagged ezrin or anillin mutants with RFP-α-tubulin. Anaphase onset was set as 0 min. The arrow indicates irregular waviness of the equatorial cortex. (**C**, **D**) Time courses of cleavage furrow widths (C) and maximum furrowing rate (D) in live-cell imaging in B. Mean ± SE of at least 10 cells from at least two independent experiments for each condition (* *p* < 0.05, ** *p* < 0.01, two-tailed *t*-test). **(E**) The frequency of cells containing the cleavage furrow with irregular waviness. Time frames of live images of anillin-depleted cells expressing EGFP-anillin or ezrin-C-term-EGFP-anillin-C-term, at which their furrow widths just reached < 10 μm, were selected for the analysis. At least 14 cells pooled from three independent experiments were analyzed.

The drastic restoration of furrow ingression by ezrin-C-term-EGFP-anillin-C-term in endogenous anillin-depleted cells prompted us to investigate whether the chimeric gene can support cell proliferation in the absence of endogenous anillin. For this, we established a HeLa cell line stably expressing ezrin-C-term-EGFP-anillin-C-term. In WT cells, anillin depletion for 4 d resulted in a drastic decrease in cell proliferation compared to mock-depleted control in a colorimetric assay, accompanying drastic multinucleation (Fig. 5A-C). On the other hand, the proliferation of ezrin-C-term-EGFP-anillin-C-term-expressing cells was not affected by the depletion of endogenous anillin (Fig. 5B). Consistent with this, ezrin-C-term-EGFP-anillin-C-term substantially suppressed multinucleation upon anillin depletion (Fig. 5C). Therefore, ezrin-C-term-EGFP-anillin-C-term could functionally substitute for endogenous anillin in supporting cell viability and proliferation.

**Fig. 5:**
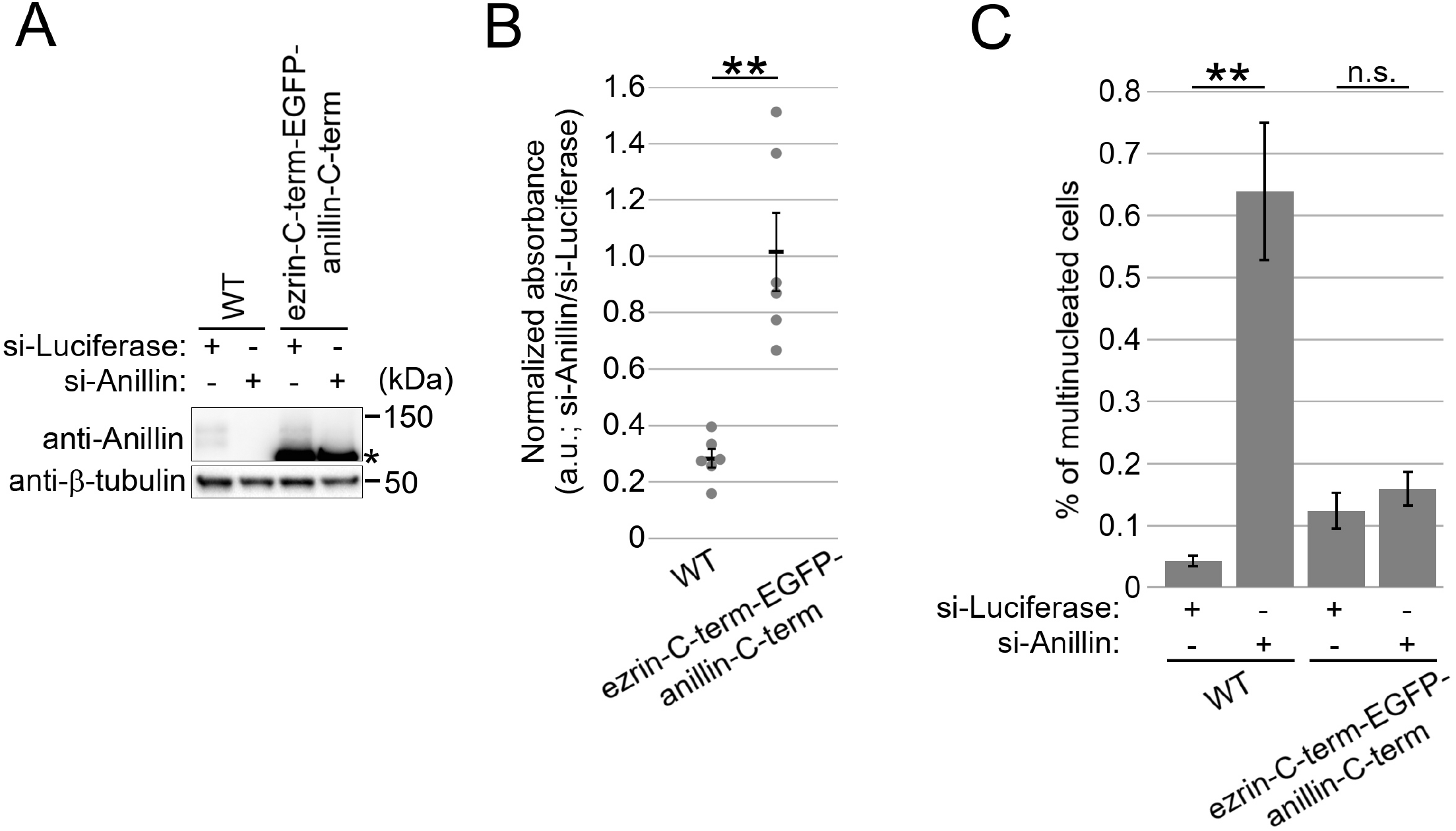
Ezrin C-terminal domain can substitute for anillin actin- and myosin-binding domains for supporting cell proliferation. **(A)** Immunoblotting of anillin in mock- or anillin-depleted WT and ezrin-C-term-EGFP-anillin-C-term HeLa cells. β-tubulin was detected as a loading control. The asterisk indicates a cross-reaction of anillin antibody to the chimeric protein. **(B)** Colorimetric cell proliferation assay in mock- or anillin-depleted WT and ezrin-C-term-EGFP-anillin-C-term HeLa cells. Absorbance in anillin-depleted samples was normalized to that of corresponding mock-depleted controls. Mean ± SE of 6 samples from 3 independent experiments. **(C)** Frequency of multinucleation in B. Mean ± SE of three independent experiments. At least 258 cells were analysed for each condition. (** *p* < 0.01, two-tailed *t*-test).

### Incompatibility of anillin domains in supporting ezrin’s role in cell division

To further understand the exchangeability of the actin-binding or membrane-associated domains between ezrin and anillin, we tested whether the chimeric proteins substitute for ezrin in regulating cortical dynamics during cytokinesis. For this, we co-depleted ezrin, moesin, and radixin in HeLa cells with or without the stable exogenous expression of ezrin mutants and chimeras (Fig. 6A). During cytokinesis in animal cells, membrane blebs frequently formed at the cell poles, presumably for pressure release [31]. In mock-depleted control cells, the formation of blebs was immediately followed by subsequent retraction, limiting the expansion of bleb structures (Fig. 6B, C, and Movie 1). Co-depletion of ERM proteins in WT HeLa cells did not affect furrow ingression but frequently caused abnormally large blebs over 40 μm^2^ at the cell poles (Fig. 6B, C, and Movie 2). The drastic increase in bleb size indicates severe defects in the bleb retraction process by depletion of ERM proteins. Similar disorganization of the polar cortex has been reported in *Drosophila* cells depleted of moesin, the only ERM protein in that organism [17]. The expression of ezrin-EGFP drastically suppressed the formation of the abnormally large blebs in ERM-depleted cells (Fig. 6B, C, and Movie 3). This result suggests that ezrin sufficiently mediates bleb retraction in the absence of radixin and moesin. Next, we tested the effects of the expression of ezrin-FERM-EGFP-anillin-MABD or ezrin-C-term-EGFP-anillin-C-term on the blebbing dynamics in ERM-depleted cells. Though expression levels of these chimeric proteins were relatively low (Fig. S1), both of them substantially accumulated to cortical blebs (Fig, 6A, B, and Movie 4 and 5). This is consistent with the fact that both the FERM and anillin-C-term domains have abilities to localize to cortical blebs [32, 33]. However, both chimera proteins failed to restore the regular blebbing dynamics in ERM-depleted cells (Fig. 6B and C). These results demonstrate that both the actin- and membrane-associated domains of anillin are not compatible with the corresponding domains of ezrin in the process.

**Fig. 6:**
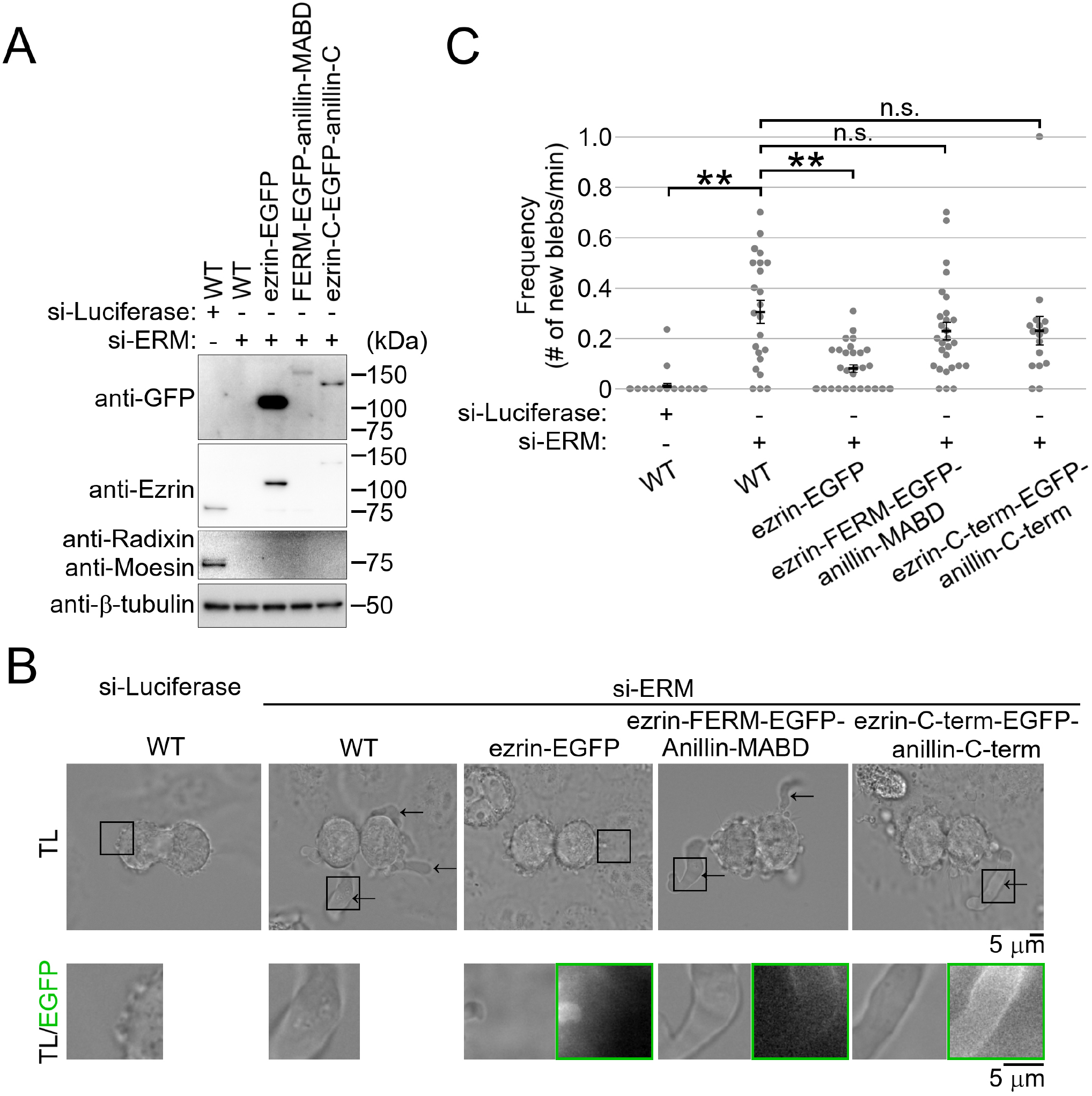
Effects of the expression of the chimeric proteins on polar cortical blebbing in ERM-depleted cells. (**A**) Immunoblotting of ERM proteins and GFP in mock- or ERM-depleted cells expressing EGFP-tagged exogenous genes. β-tubulin was detected as a loading control. (**B**) Live images of mock- or ERM-depleted HeLa cells expressing ezrin-EGFP or chimeric proteins in cytokinesis. The arrows indicate abnormally large blebs. Polar blebs are 3× enlarged in the bottom panels. Transmitted light (TL) and fluorescence (EGFP) microscopies are shown. **(C**) The frequency of abnormally large bleb formation. Mean ± SE of at least 16 cells from at least four independent experiments (** *p* < 0.01, two-tailed *t*-test).

## Discussion

In this study, we investigated the molecular basis of the semi-compatibility between ERM proteins and other actin-membrane associated proteins in cytokinesis. First, we addressed the mechanism underlying the compensatory over-accumulation of ezrin upon depletion of other actin-membrane-associated proteins. Previous studies showed that the drastic changes in the dynamics of cortical association/dissociation of ERM proteins accompany the control of their localization [30, 34]. However, the FRAP profile of ezrin-EGFP did not change upon compensatory over-accumulation in supervillin- or anillin-depleted cells, indicating that this process takes place without the active regulation of ezrin’s turnover dynamics at the cortex. Meanwhile, the selective over-accumulation of ezrin FERM or the C-term at the furrow took place upon depletion of anillin or supervillin, respectively, indicating that these proteins affect the localization of ezrin through different mechanisms. A possible interpretation of these results is that ezrin competes for limited binding sites at the actin cytoskeleton or membrane with supervillin or anillin, respectively. This idea is at least consistent with the fact that both ezrin and anillin localized to the plasma membrane surface through their interaction with PIP2 [3, 35–38]. Another possibility is that the absence of anillin or supervillin may change the actin or membrane scaffolds’s states so that ezrin domains can preferentially associate with them. The extent of the compensatory over-accumulation was much smaller for the ezrin domain truncates than for full-length ezrin, suggesting that both the FERM and C-term are required for the maximum association of ezrin to the division site. These results possibly explain why ERM proteins became engaged in furrowing activity only when both anillin and supervillin were co-depleted in our previous study [15]; only in this condition, ezrin may get full access to the binding sites both at actin cytoskeleton and membrane that are responsible for furrow ingression.

Ezrin-FERM-EGFP-anillin-MABD could not substitute for endogenous anillin in supporting furrow ingression. Therefore, despite the possibility that ezrin and anillin share their binding sites at the membrane, their membrane-associated domains are not functionally compatible. In contrast, the ezrin C-term could sufficiently substitute for anillin MABD in supporting furrow ingression and long-term cell proliferation. This result demonstrates the potential of ezrin C-term to associate with the essential fraction of the actin cytoskeleton for cytokinesis progression. It also suggests the flexibility in the choice of actin-binding moiety for supporting anillin’s essential role in cytokinesis, which is consistent with a previous report [39]. However, the frequent membrane deformation in the furrow ingression supported by ezrin-C-term-EGFP-anillin-C-term indicates the importance of the unique properties of anillin MABD [19, 20] in achieving efficient and smooth membrane invagination at the edge of the cleavage furrow. On the other hand, either anillin-C-term or anillin MABD could not substitute for the corresponding domain of ezrin in supporting the regular dynamics of bleb retraction, highlighting poor compatibility between ezrin and anillin in the process. Extra-long blebs form through sequential generations of secondary blebs, presumably when tethering of the actin cytoskeleton to the membrane at the blebs is not strong enough to resist inner cytoplasmic pressure [31]. Therefore, these chimeric proteins may not support the formation of a rigid actin-membrane tether. Ultrastructural studies have revealed that a relatively isotropic cage-like 3D network of the actin cytoskeleton filling the entire volume of bleb forms during the bleb retraction [32, 40]. On the other hand, the actin cytoskeleton with the contractile ring is packed more tightly, forming an anisotropic purse-string like meshwork during furrow ingression in different organisms [41–43]. Such polymorphism in actin cytoskeletal ultrastructure in different cellular processes may set a stringent limit to the functional compatibility among different actin-binding moieties. Our results shed light on diverse modes of actin-membrane interactions supported by different actin-membrane associated proteins, enabling complex regulation of cell deformation during cytokinesis.

---

**Fig. S1:**
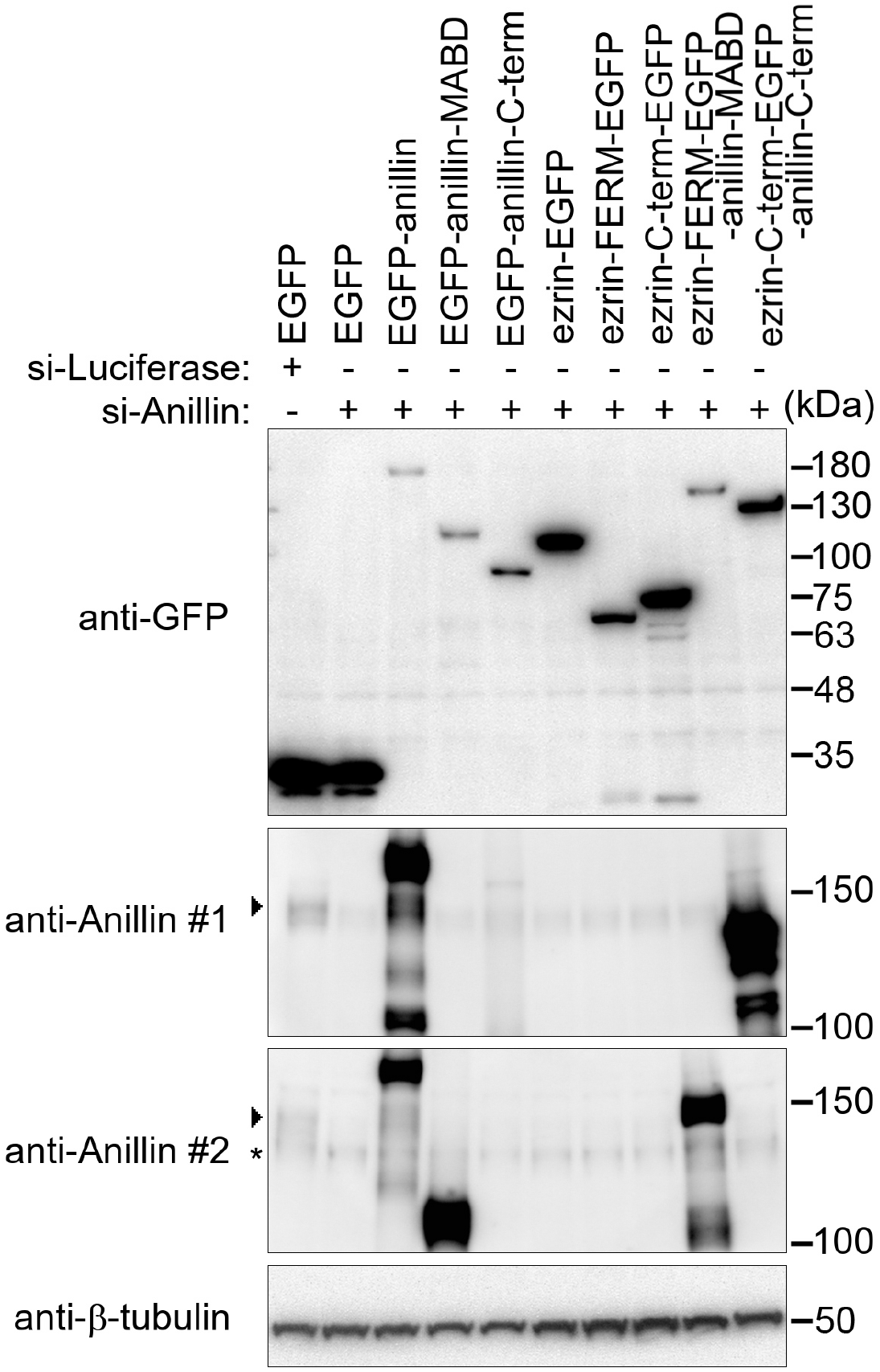
Immunoblotting of the cells expressing EGFP-tagged exogenous genes. Immunoblotting of GFP and anillin in the cells expressing EGFP-tagged exogenous genes. Anillin was detected with two different antibodies to confirm depletion of endogenous anillin in the cells expressing different chimeric proteins. The arrowheads indicate endogenous anillin and the asterisk indicates nonspecific bands. β-tubulin was detected as a loading control.

**Fig. S2:**
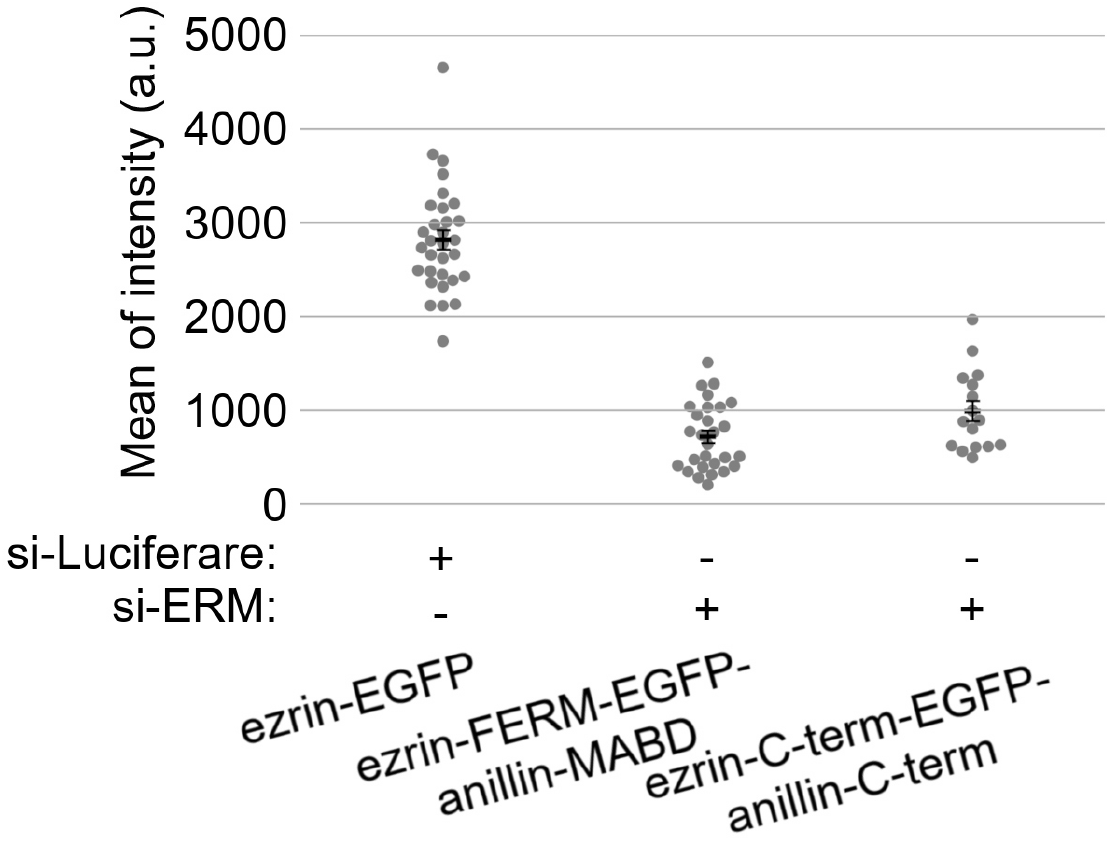
Expression level of EGFP-tagged proteins. The whole-cell fluorescence intensity of EGFP-tagged proteins in the cells analysed in Fig. 6C. Mean fluorescence intensity in the whole cell area was subtracted by background fluorescence intensity outside the cells. Mean ± SE of at least 16 cells from at least four independent experiments.

**Movie 1: A mock-depleted WT HeLa cell undergoing cytokinesis**

Left: A transmitted light (TL) microscopy. Right: A fluorescent microscopy (GFP channel). Movie is shown at 300× real time. Field of view is 50 μm × 40 μm for each channel.

**Movie 2: An ERM-depleted WT HeLa cell undergoing cytokinesis**

Left: A TL microscopy. Right: A fluorescent microscopy (GFP channel). Movie is shown at 300× real time. Field of view is 70 μm × 50 μm for each channel.

**Movie 3: An ERM-depleted ezrin-EGFP HeLa cell undergoing cytokinesis**

Left: A TL microscopy. Right: A fluorescent microscopy (GFP channel). Movie is shown at 300× real time. Field of view is 60 μm × 50 μm for each channel.

**Movie 4: An ERM-depleted ezrin-FERM-EGFP-anillin-MABD HeLa cell undergoing cytokinesis**

Left: A TL microscopy. Right: A fluorescent microscopy (GFP channel). Movie is shown at 300× real time. Field of view is 60 μm × 60 μm for each channel.

**Movie 5: An ERM-depleted ezrin-C-term-EGFP-anillin-C-term HeLa cell undergoing cytokinesis**

Left: A TL microscopy. Right: A fluorescent microscopy (GFP channel). Movie is shown at 300× real time. Field of view is 80 μm × 50 μm for each channel.

## Acknowledgement

We thank Sarada Bulchand for commenting on the draft, the Open Facility, Global Facility Center, Creative Research Institution, Hokkaido University for allowing us to use their instrument. This work was supported by the Nomura Gakugei Foundation and the Sasakawa Scientific Research Grant from The Japan Science Society to S.H., Grants-in-Aid for Scientific Research C (18K06201) to A.K., Grant-in-Aid for Scientific Research on Innovative Areas “Information physics of living matters” (20H05522) to M.K., DBT/Wellcome Trust India Alliance (India Alliance; IA/I/14/1/501317) to M.M., Grants-in-Aid for Scientific Research B (19H03219), on Innovative Areas “Singularity Biology (No.8007)” (19H05413), Fostering Joint International Research B (19KK0181), and JSPS Bilateral Joint Research Project (JPJSBP120193801) of MEXT, the Princess Takamatsu Cancer Research Fund, the Kato Memorial Bioscience Foundation, the Orange Foundation, the Smoking Research Foundation, the Suhara Memorial Foundation, and the Nakatani Foundation to R.U.

## Author Contributions

Conceptualization, G.Y., S.H., and R.U.; Methodology, A.K., M.K., M.M., and R.U.; Investigation, G.Y., S.H., and R.U.; Formal Analysis, G.Y., S.H., and R.U.; Resources, A.K., M.K., M.M., and R.U.; Writing – Original Draft, R.U.; Writing – Review & Editing, G.Y., S.H., A.K., M.M., and R.U.; Funding Acquisition, S.H., A.K., M.K., M.M., and R.U.

