## Supplementary material for "Molecular basis of functional compatibility between ezrin and other actin-membrane associated proteins during cytokinesis": TableS1

Table S1: List of the plasmids used in this study

| Plasmid ID | Plasmid name | Insert (GenBank accession #) | Detailed description |
| --- | --- | --- | --- |
| N/A | pPB-EF1a-MCS-EGFP | N/A | MCS-EGFP cassette of the pEGFP-N1 vector was inserted into NheI/NotI sites of a PiggyBac-based vector (SBI). |
| N/A | pPB-EF1a-EGFP-MCS | N/A | EGFP-MCS cassette of the pEGFP-C1 vector was inserted into NheI/BamHI sites of a PiggyBac-based vector (SBI). |
| pSH35 | ezrin-EGFP | Ezrin, full length (NM_001111077.1) | The cDNA fragment was amplified by PCR using primers;  5′- ATAAAGCTTCACCATGCCGAAACCAATCAATGTCCGAG-3′ and 5′-ATAGTCGACTGCAGGGCCTCGAACTCGTCGATGC-3′, once cloned into HindIII/SalI sites of pEGFP-N1, and then into XhoI/AgeI sites of pPB-EF1a-MCS-EGFP. |
| pSH36 | ezrin-FERM-EGFP | Ezrin truncate (NM_001111077.1) 1-309 a.a. | The cDNA fragment was amplified by PCR using primers;  5′- ATAAAGCTTCACCATGCCGAAACCAATCAATGTCCGAG -3′ and 5′- ATAGTCGACTGGGCCTGGGCCTTCATCTGCTGCACC -3′, once cloned into HindIII/SalI sites of pEGFP-N1, and then into XhoI/AgeI sites of pPB-EF1a-MCS-EGFP. |
| pSH37 | ezrin-C-term-EGFP | Ezrin truncate (NM_001111077.1) 296-586 a.a. | The cDNA fragment was amplified by PCR using primers;  5′- ATAAAGCTTCACCATGAAGCCTGACACCATCGAGGTGC -3′  and 5′- ATAGTCGACTGCAGGGCCTCGAACTCGTCGATGC -3′, once cloned into HindIII/SalI sites of pEGFP-N1, and then into XhoI/AgeI sites of pPB-EF1a-MCS-EGFP. |
| pSH41 | EGFP-anillin-MABD | Anillin truncate  (NM_018685.5)  1-606 a.a. | The cDNA fragment was amplified by PCR using primers;  5′- ATACTCGAGAAATGGATCCGTTTACGGAGAAACTGCTGG -3′ and 5′-ATAGAATTCTTATGAGGAGATATTCAGTGCATCTTCCTGTTCTTCGC -3′, and cloned into XhoI/EcoRI sites of pPB-EF1a-EGFP-MCS. |
| pSH42 | ezrin-FERM-EGFP-anillin-MABD | N/A | EGFP-Anillin-MABD fragment from pSH41 was cloned into AgeI/NotI sites of pSH36. |
| pSH44 | EGFP-anillin-C-term | Anillin truncate  (NM_018685.5)  609-1124 a.a  *corresponding to NM_001284301.2, 572-1087 a.a. | The cDNA fragment was amplified by PCR using primers;  5′-ATAGAGCTCAAATGTTACTTGCACCATTGGCACAAACAG -3′ and 5′- ATACCGCGGTTAAGGCTTTCCAATAGGTTTGTAGCAAGC -3′, once cloned into SacI/SacII sites of pEGFP-C1 vector, and then into AgeI/SacII sites of pPB-EF1a-EGFP-MCS. |
| pSH43 | ezrin-C-term-EGFP-anillin-C-term | N/A | The EGFP-Anillin-C-term fragment was digested from pEGFP-anillin-C-term (the precursor of pSH44; *see above*) using AgeI/PspOMI, and cloned into AgeI/NotI sites of pSH37. |
| pSH48 | EGFP-anillin | Anillin, full length  (NM_018685.5) | Full-length anillin was amplified by PCR using primers;  5′- ATAGAGCTCAAATGGATCCGTTTACGGAGAAACTGCTGG -3′ and 5′- ATACCGCGGTTAAGGCTTTCCAATAGGTTTGTAGCAAGC -3′, digested using KpnI/SacII. This resulted in an anillin fragment whose N-terminal end is removed by the digestion of the endogenous KpnI site (fragment #1). The anillin-N-term fragment was digested from pSH41 using XhoI/KpnI (fragment#2). Then, fragment#1 and #2 were cloned into XhoI/SacII sites of pPB-EF1a-EGFP-MCS. |
| pSH49 | ezrin-EGFP  (ezrin siRNA- resistant) | Ezrin, full length (NM_001111077.1) | An ezrin-N-terminal fragment was digested from pEGFP-N1-ezrin (the precursor of pSH35; *see above*) using HindIII/SphI (fragment #3). Then an ezrin cDNA fragment containing siRNA-resistant sequence was amplified by PCR using primers;  5′- ATAGCATGCGGAACACAGAGGAATGTTAAAGGACAATGCTATGTTGGAATACCTGAAGATTGCTCAGG -3′ and 5′-ATAGTCGACTGCAGGGCCTCGAACTCGTCGATGC-3′, and digested using SphI/SalI (fragment #4). Fragment #3 and #4 were once cloned into HindIII/SalI sites of pEGFP-N1, and then into XhoI/AgeI sites of pPB-EF1a-MCS-EGFP. |
| pSH50 | ezrin-FERM-EGFP (ezrin siRNA- resistant) | Ezrin truncate (NM_001111077.1) 1-309 a.a. | An ezrin cDNA fragment containing siRNA-resistant sequence was amplified by PCR using primers;  5′- ATAGCATGCGGAACACAGAGGAATGTTAAAGGACAATGCTATGTTGGAATACCTGAAGATTGCTCAGG -3′ and 5′- ATAGTCGACTGGGCCTGGGCCTTCATCTGCTGCACC -3′, and digested using SphI/SalI (fragment #5). Fragment #3 (prepared for pSH49; *see above*) and #5 were once cloned into HindIII/SalI sites of pEGFP-N1, and then into XhoI/AgeI sites of pPB-EF1a-MCS-EGFP. |
| pSH52 | ezrin-FERM-EGFP-anillin-MABD (ezrin　siRNA-resistant) | N/A | EGFP-Anillin-MABD fragment from pSH41 was cloned into AgeI/NotI sites of pSH50. |
